# Genetic Basis of Transcriptome Diversity in *Drosophila melanogaster*

**DOI:** 10.1101/018325

**Authors:** Wen Huang, Mary Anna Carbone, Michael M. Magwire, Jason A. Peiffer, Richard F. Lyman, Eric A. Stone, Robert R. H. Anholt, Trudy F. C. Mackay

## Abstract

Understanding how DNA sequence variation is translated into variation for complex phenotypes has remained elusive, but is essential for predicting adaptive evolution, selecting agriculturally important animals and crops, and personalized medicine. Here, we quantified genome-wide variation in gene expression in the sequenced inbred lines of the *Drosophila melanogaster* Genetic Reference Panel (DGRP). We found that a substantial fraction of the *Drosophila* transcriptome is genetically variable and organized into modules of genetically correlated transcripts, which provide functional context for newly identified transcribed regions. We identified regulatory variants for the mean and variance of gene expression, the latter of which could often be explained by an epistatic model. Expression quantitative trait loci for the mean, but not the variance, of gene expression were concentrated near genes. This comprehensive characterization of population scale diversity of transcriptomes and its genetic basis in the DGRP is critically important for a systems understanding of quantitative trait variation.

## Introduction

Genetic variation for quantitative traits is a universal property of evolving populations. Elucidating the general principles that underlie the genotype-phenotype map is critical for understanding natural selection and evolution, improving the efficacy of animal and plant breeding, and identifying targets for treating human diseases. Numerous quantitative trait loci (QTLs) have been identified in linkage and association mapping populations by scanning polymorphic markers across the genome. However, QTLs rarely map to genes or causal genetic variants and typically account for only a small fraction of total genetic variation (Flint and Mackay 2009; Manolio et al. 2009). This makes interpreting the functional roles of QTLs and dissecting the genetic architecture of quantitative traits particularly challenging.

By extension of the central dogma of molecular biology, it is generally accepted that a QTL generates phenotypic variation by introducing variation in protein sequence and/or abundance of gene products (Mackay et al. 2009). Variation in the abundance of gene products constitutes an important class of quantitative traits and can be measured with great precision and high throughput. This provides the opportunity to identify expression QTLs (eQTLs) that control variation in global mRNA levels. Furthermore, while the relative importance of structural and regulatory variation remains debatable, mounting evidence has indicated that regulatory variation could be a significant source of phenotypic variation. In particular, there is increasing evidence that QTLs associated with organismal phenotypes are more likely to be eQTLs than other variants of similar allele frequencies in the genome (Nicolae et al. 2010).

Genetic studies of global gene expression in model organisms and human populations have found that a substantial fraction of gene expression traits is heritable (e.g. Brem et al. 2002; Cheung et al. 2003; Schadt et al. 2003; Ayroles et al. 2009). While both local (*cis*) and distal (*trans*) eQTLs have been detected, in most cases eQTLs near genes tend to be more common and have larger effects. Conventionally, individuals within each genotype class of an eQTL share the same mean of expression, which differs among individuals of different genotypes (we call these mean eQTLs or simply eQTLs throughout this study). More recently, another class of QTLs for which there is a difference in the variance of phenotypes between individuals with different genotypes has been identified for both gene expression (Hulse and Cai 2013; Brown et al. 2014) and organismal phenotypes (Rönnegård and Valdar, 2011; Shen et al. 2012; Yang et al. 2012). These variance QTLs are of interest because differences in the variance of gene expression among different genotypes at a focal locus can be induced by epistasis between the focal locus and one or more interacting loci (Rönnegård and Valdar, 2011), thereby providing a simple approach for identifying QTLs participating in genetic interactions (Brown et al. 2014).

The *Drosophila melanogaster* Genetic Reference Panel (DGRP) consists of 205 inbred lines with whole genome sequences (Mackay et al. 2012; Huang et al. 2014). The DGRP harbors molecular variation for more than four million loci (∼ one every 50 base pairs) and exhibits quantitative genetic variation for many organismal phenotypes (Huang et al. 2014 and references therein), facilitating genome-wide association (GWA) mapping in a scenario where nearly all variants are known. Recent GWA studies in the DGRP indicate that the inheritance of most organismal quantitative traits in *Drosophila* is complex, involving many genes with small additive effects as well as epistatic interactions (Mackay et al. 2012; Huang et al. 2012; Swarup et al. 2013; Mackay, 2014).

A small-scale study of 40 DGRP lines has previously revealed substantial quantitative genetic variation in gene expression in the DGRP (Ayroles et al. 2009). The genetically variable transcripts cluster into modules of highly correlated expression traits associated with distinct biological processes (Ayroles et al. 2009). More recently, an eQTL mapping analysis in this subset of DGRP lines has identified *cis* eQTLs within 10kb of more than 2,000 genes (Massouras et al. 2012).

As QTL mapping studies in the DGRP accumulate information on the genetic basis of many organismal traits, a comprehensive characterization of the diversity of transcriptomes and its genetic basis in the entire DGRP becomes critically important. In the present study, we identify unannotated transcriptional units in the *Drosophila* genome using RNA-Seq and quantify gene expression using genome tiling microarrays. We then comprehensively characterize the genetic diversity of gene expression in the DGRP. Finally, we identify eQTLs that control the mean and variance of global gene expression, and show that the latter can frequently be explained by interactions with *cis*-eQTLs.

## Results

### Identification of novel transcribed regions

Recent efforts to characterize genome-wide transcription in human cells found that approximately three quarters of the human genome is transcribed into primary transcripts and more than 60% of the genomic bases represent processed mature RNA transcripts (Djebali et al. 2012). Pervasive transcription appears to be a common feature for eukaryotic genomes (Dinger et al. 2009). Approximately 75% of the *D. melanogaster* genome is transcribed at least temporarily during development, and thousands of novel transcribed regions have been identified, the majority of which do not appear to code for proteins (Graveley et al. 2011). With the exception of a small number of long non-coding RNAs (lncRNAs) in mammals whose regulatory roles are well established (Lee 2012), the functional implications of pervasive transcription and non-coding RNAs (ncRNAs) remain to be resolved.

The DGRP provides a platform to study the molecular quantitative genetics and functions of RNAs by associating them with genetic determinants of gene expression and expression of other RNAs. To identify unannotated novel transcribed regions (NTRs), we sequenced poly (A)+ RNAs of adult flies pooled from 192 DGRP lines using 100 bp paired-end sequencing, separately for females and males. Approximately 100 M cDNA fragments were sequenced in both sexes (Table S1). We aligned the sequence reads to the annotated transcriptome and reference genome and used the resulting overlapping alignments to assemble transcript models. Approximately 4.5% (females) and 6.7% (males) of mapped reads do not overlap with any annotated exons and may represent unannotated transcriptional units (Table S1). We merged overlapping transcript models in females and males and compared them with the FlyBase (Release 5.49) annotation to identify NTRs. We found 1,669 and 2,192 transcripts derived from 1,628 intronic and 1,876 intergenic regions, respectively, representing a total of 3.6 M unannotated bases in processed RNAs – an approximately 11% addition to the existing annotations. In addition, a total of 2,807 novel alternatively spliced isoforms were found for 2,049 annotated genes. We characterized NTRs for the size of processed transcripts they produce, nucleotide composition, sequence conservation, and propensity to harbor polymorphic DNA variants. NTRs do not differ qualitatively from annotated ncRNAs. Compared to protein-coding genes, both NTRs and annotated ncRNAs have shorter transcripts, lower GC content, weaker sequence conservation and slightly higher density of DNA variants (Figure S1).

We estimated the expression of annotated genes and NTRs in the pooled samples of females and males. Not surprisingly, NTRs are generally expressed at a much lower level than annotated genes (Figure 1); more highly and ubiquitously expressed genes were more likely to be detected by previous annotation efforts. We reasoned that spurious non-functional NTRs identified in RNA-Seq would not be genetically variable in subsequent quantitative genetic analyses using an independent expression platform. Therefore, we did not filter NTRs by their expression level, a common practice to eliminate erroneous transcript reconstruction in RNA-Seq.

**Figure 1:**
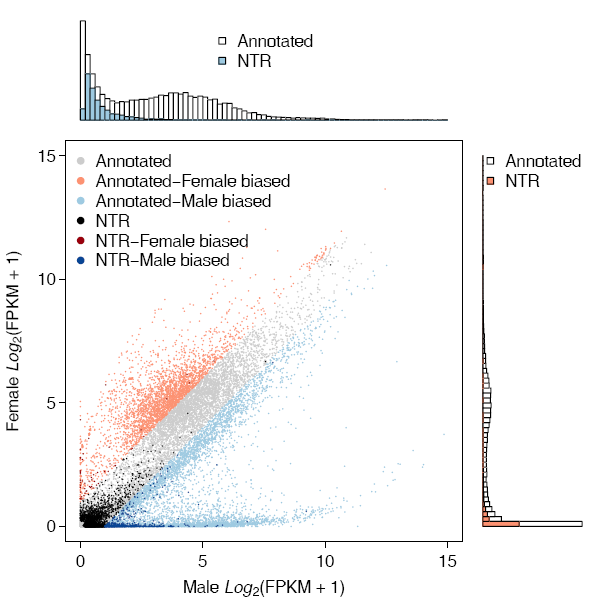
RNA-Seq in the DGRP reveals many novel transcribed regions. The scatter plot compares gene expression of annotated genes and NTRs in females and males. Genes with expression differences of two-fold or more between the sexes are considered to have sex-biased expression. The histograms depict the distribution of gene expression in females (left) and males (top), with colored bars showing the distributions for NTRs.

### Transcriptome diversity in the DGRP

We used Affymetrix *Drosophila* 2.0 genome tiling arrays to measure expression of annotated genes and NTRs in 185 DGRP lines, with two biological replicates for each sex. We estimated the overall expression of genes by median polish of background corrected and quantile normalized probe expression. Only probes which uniquely and entirely map to constitutive exons and do not contain common (non-reference allele frequency > 0.05) variants were used.

We used a linear mixed model to test for the effect of sex (sexual dimorphism) and partition the variance in gene expression into three sources, including between-line (genetic) variance, variance in sex by line interaction (genetic variance in sexual dimorphism), and within-line (environmental) variance. As expected, given that sexual dimorphism is common for *D. melanogaster* gene expression traits (Ranz et al. 2003; Parisi et al. 2004; Ayroles et al. 2009), the vast majority (16,445/18,140, 90.6%) of genes showed significant mean differences (FDR < 0.05) between females and males, including NTRs, of which 80.9% show sex-biased expression (2,743/3,391) (Table S2). Gene set enrichment analysis revealed that genes with female biased expression were enriched for several biological processes primarily associated with DNA replication, DNA repair and the cell cycle, while genes with male biased expression were enriched for genes involved in reproduction (Table S3, Figure S2). Furthermore, genes with sex-biased expression are highly enriched for ovary- and testis-specific genes, respectively (Figure S3). A substantial fraction of genes (2,388/18,140, 13.2%, of which 106/3,391 (3.1%) were NTRs) show significant (FDR < 0.05) sex by line interaction, indicating that the degree of sexual dimorphism as a quantitative trait is genetically variable for these genes (Table S2). The lower proportion of NTRs showing sexual dimorphism and sex by line interaction is likely a result of their low expression and thus smaller effects. Because of the widespread sexual dimorphism and sex by line interaction, we performed all subsequent analyses in females and males separately.

We next asked to what extent variation in gene expression is heritable. We tested the significance of the among-line variance component and estimated the broad sense heritability (*H*^2^) for each gene expression trait as the proportion of total variance explained by between-line differences. Among the 18,140 annotated genes and NTRs, a total of 7,626 unique genes showed significant (FDR < 0.05) genetic variability in expression in either sex (Table S4). Among these genetically variable transcripts, 4,308 had a significant genetic component in females, 5,814 in males, and 2,496 in both sexes (Table S4, Figure 2A-B). Remarkably, 231 and 430 NTRs are genetically variable in females and males, respectively, and 111 NTRs are genetically variable in both sexes (Table S4). Estimates of broad sense heritability for genes with heritable variation in expression range from 0.034 – 0.946 in both sexes (Figure 2).

**Figure 2:**
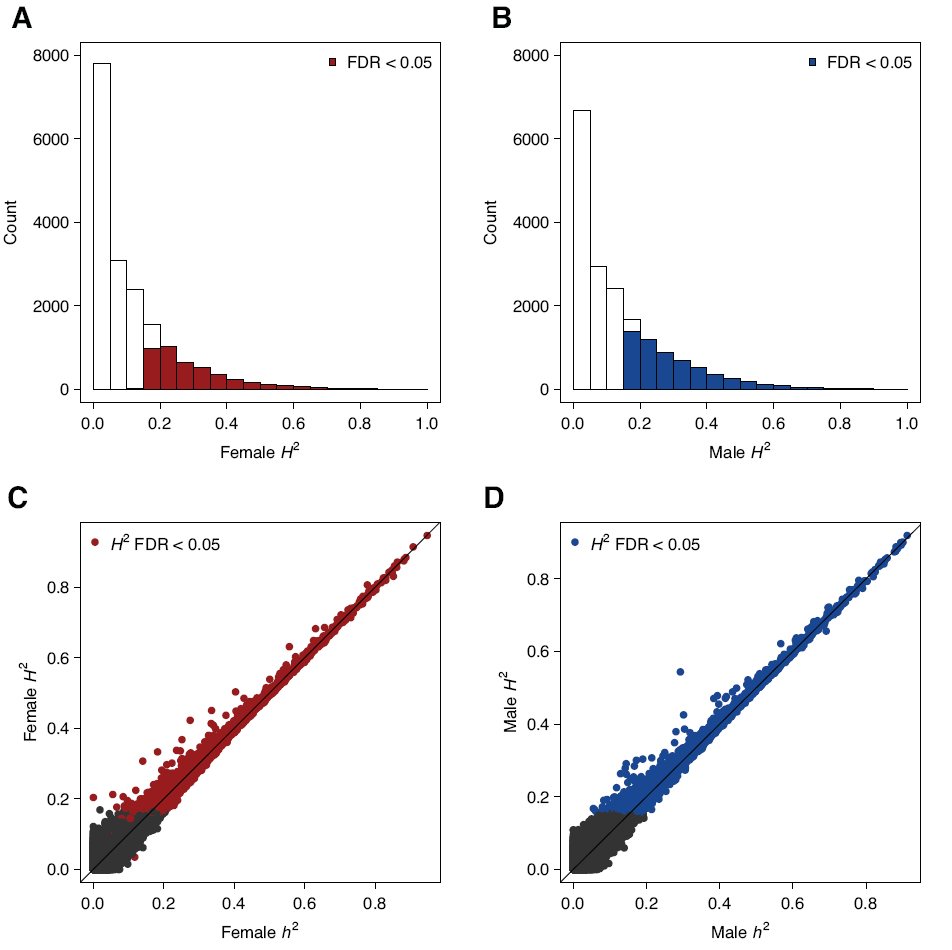
Genetic variation of gene expression. (**A** and **B**) Distribution of broad sense heritability (*H*^2^) for gene expression traits. (A) Females. (B) Males. (**C** and **D**) Relationship between narrow sense heritability (*h*^2^) and *H*^2^ for gene expression traits. (C) Females. (D) Males. Genetically variable genes (FDR < 0.05) are color-coded as indicated.

Given the availability of complete genome sequences, we can compute the genetic covariance among the DGRP lines, which measures the genetic similarity between pairs of lines assuming an infinitesimal model. This allows us to estimate the proportion of phenotypic variance in gene expression explained by the additive genetic variance (or narrow sense heritability, *h*^2^) using a mixed effects model (Yang et al. 2010; Ober et al., 2012). Interestingly, *h*^2^ of the vast majority of genes captures most of the total genetic variance (Table S4, Figure 2C-D). While large differences between *h*^2^ and *H*^2^ indicate a large contribution of non-additive gene action (i.e. dominance and/or epistasis), the opposite is not necessarily true (Mackay 2014). Epistatic gene action can lead to largely additive variance if the minor allele frequencies (MAF) of interacting loci are low (Hill et al. 2008; Mackay 2014).

Among the 185 DGRP lines, 99 were infected with the endosymbiotic bacterium *Wolbachia pipientis* (Huang et al. 2014). We tested the effect of *Wolbachia* infection on gene expression, conditional on five polymorphic major inversions and the first ten principal components (PCs) of common variants. Because lines are nested within the *Wolbachia* infection, it is not possible to separate the *Wolbachia* effect from between-line variation. Nevertheless, by accounting for major inversions and top genotypic PCs, we aim to test for the effect of *Wolbachia* independent of genetic differentiation between the lines. Overall, *Wolbachia* infection has only minor effects on gene expression, and the effects are female-specific (Figure S4). In particular, genes that are down-regulated in lines positive for *Wolbachia* are largely ovary-specific (Figure S4).

Many large chromosomal inversions are polymorphic in the DGRP, some of which are at relatively high frequency. We tested the effects of each of the five major segregating inversions on the expression of genetically variable transcripts. For each inversion, we grouped lines segregating for the inversion into a third genotype class in addition to the two inbred genotypes, noting that frequencies of inversions within these lines may vary. At FDR < 0.05, there are 125 (20), 9 (13), 35 (26), 17 (32), 21 (39) genes in females (males) whose expression is affected by *In(2L)t*, *In(2R)NS*, *In(3R)P*, *In(3R)K*, and *In(3R)Mo*, respectively (Table S5). We also tested whether inversions preferentially affect expression of genes within the inverted regions. Such a local effect could be indicative of the accumulation of *cis*-regulatory mutations after the inversions arose in the population. Interestingly, *In(2L)t* and *In(3R)Mo* preferentially affect genes within the boundaries of their respective inversions, in both sexes, while other inversions do not appear to do so (Figure S5).

### Modules of genetically correlated transcripts

We have shown previously using 40 DGRP lines that genetically variable transcripts are not independent, but cluster into a smaller number of genetically correlated co-expression modules whose members often contribute to the same biological processes (Ayroles et al. 2009). To expand the investigation to the entire DGRP, we first estimated the genetic component of expression for genetically variable transcripts after adjusting for *Wolbachia* infection status. Because inversions affect the expression of only a small number of genes (Figure S5) and they are genuine genetic effects, we did not adjust for their effects in our analysis of correlated gene expression.

We used Modulated Modularity Clustering (MMC; Ayroles et al. 2009; Stone and Ayroles, 2009) to identify clusters of genetically correlated genes. This algorithm derives modules such that the absolute value of the genetic correlation among transcripts is maximized within modules and minimized between modules. We found a few large modules of high connectivity in both sexes (Figure 3; Table S6). These modules are not merely statistical constructs, but are frequently enriched for genes within the same gene ontology terms (Tables S7, S8; Figure S6), indicating that genes with genetically correlated transcripts tend to fall within the same biological pathways. Indeed, the genetic correlation in expression of genes belonging to the same GO pathways is significantly higher than that between genes in different GO pathways (Figures S7, S8). Therefore, functions of computationally predicted genes within these modules can be inferred with functional annotations of other genes in the module using the principle of ‘guilt by association’ (Ayroles et al. 2011).

**Figure 3:**
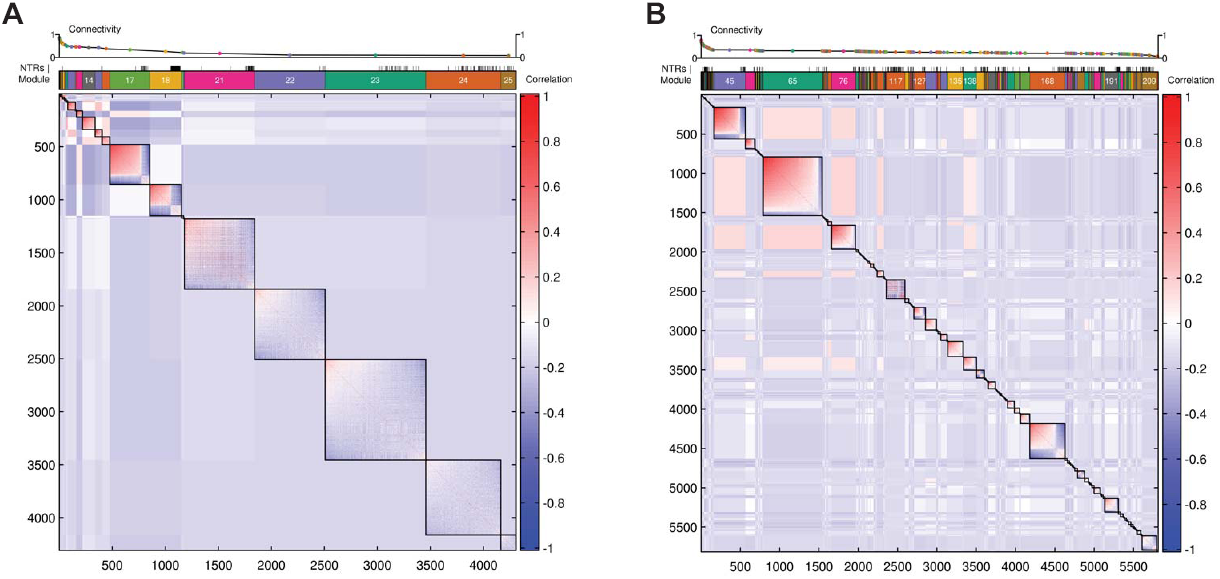
Genetically correlated modules of gene expression traits. (**A** and **B**) Heat maps from MMC analyses. Genetically variable transcripts are ordered based on their cluster membership and connectivity, which decreases from the top left corner to the bottom right corner of the heat maps. The correlation between transcripts within and between modules is depicted by the color scale bars. The modules are indicated by the colored rectangles above the heat maps, and NTRs are denoted by short vertical bars. The average connectivity within each module is given at the top of the plots. (A) Females. (B) Males.

The remaining transcripts are organized into either large modules with low connectivity (especially in females, Figure 3A) or smaller modules with relatively high connectivity (especially in males, Figure 3B). The choice between few large modules with low connectivity versus many small modules with high connectivity is affected by both the specific genetic correlation structure and the object function in the MMC clustering algorithm (Stone and Ayroles, 2009). We focused our biological inference on relatively large modules with high connectivity, which are less affected by stochastic noise in estimates of genetic correlation. Consistent with the small effect of *Wolbachia* on gene expression traits, the overall patterns of genetic correlation before and after adjusting for *Wolbachia* are largely similar (Figure S9); therefore we performed subsequent inferences based on the clustering after adjusting for *Wolbachia* infection.

Remarkably, expression of NTRs is in general negatively correlated with expression of protein-coding genes from the same expression modules – especially in males – suggesting that NTRs may act as negative regulators for expression of protein-coding genes (Figure 3, Figure S10). The mechanism by which NTRs regulate gene expression is unclear. Most NTRs, regardless of their strength of association with protein-coding genes, are distant from protein-coding genes with which they are associated (Figure S11), suggesting that NTRs function in *trans.* Among the 5,733 pairs of NTRs and protein-coding genes whose genetic correlation exceeds 0.25 in females and the 11,519 such pairs in males, only 6 and 26 had very weak homology, respectively, all of which were shorter than 30 base pairs, suggesting that NTRs do not function through base-pairing with mRNAs.

The genetic correlation in expression between NTRs and annotated genes allows us to infer putative functions of NTRs by co-expression. We used gene set enrichment analysis (GSEA) to associate (FDR < 0.05) 105 of 231 genetically variable NTRs in females and 208 of 430 genetically variable NTRs in males with at least one GO or KEGG pathway (Table S9). The majority of these associations are negative. Several pathways such as mitotic spindle organization, unfolded protein binding, and mitosis in females; and translation initiation factor activity, protein binding, and ubiquitin-protein ligase activity in males; appear to recruit a large number of NTRs (Figure S12).

### QTLs associated with mean transcript abundance

To characterize the genetic architecture of quantitative variation in gene expression, we performed GWA analyses to map expression QTLs (mean eQTLs) that regulate mean expression for all genetically variable genes. We fitted linear mixed models to adjust for *Wolbachia*, inversions and ten significant PCs of the genotypes, and estimated line means for each genetically variable transcript using best linear unbiased prediction (BLUP). The significance of association between each of the 1,913,487 individual common variants (MAF ≥ 0.05) and mean of gene expression traits was evaluated by single marker regression of the BLUP line means on marker genotypes. The empirical FDR for each gene expression trait was estimated by dividing the expected number of associations under the null hypothesis (*n* = 100 permutations) at variable *P*-value thresholds by the observed number of associations at the same *P*-value thresholds.

As expected, fewer significant eQTLs are detected as increasingly stringent FDR thresholds are applied (Table 1). By arbitrarily defining eQTLs as variants within ±1kb of the genes they influence as *cis*-eQTLs, more than 50% of genes with eQTLs have at least one *cis*-eQTL at FDR < 0.05. More *trans*-eQTLs are detected at more lenient FDR thresholds, while the increase in the number of *cis*-eQTLs is relatively small (Table 1). This result is consistent with the observation of stronger association of *cis*-eQTLs with variation in gene expression (Figure S13). At an empirical FDR < 0.20, there are 941 (females) and 1,339 (males) genetically variable gene expression traits that have at least one *cis*- and/or *trans*-eQTL, of which 31 and 114 are NTRs in females and males, respectively (Tables S10, S11). Interestingly, the proportion of genes with *cis*-eQTLs for males is substantially larger than that for females (Table 1). The association between DNA variants and gene expression is much stronger around transcription start and end sites (TSS and TES; Figure S13), where regulatory elements for transcription and RNA stability are concentrated. This observation is consistent with the distribution of *cis* eQTLs previously found in *Drosophila* and other organisms (Ronald et al. 2005; Stranger et al. 2007; Veyrieras et al. 2008; Massouras et al. 2012).

**Table 1.** Number of genes with at least one significant eQTL at different FDR thresholds.

| Sex | FDR thresholds (*cis* + *trans*)<sup>1</sup> |  |  |  |
| --- | --- | --- | --- | --- |
|  | 0.05 | 0.10 | 0.15 | 0.20 |
| Female | 503 (263 + 240) | 671 (287 + 384) | 807 (297 + 510) | 941 (308 + 633) |
| Male | 837 (533 + 304) | 1,029 (568 + 461) | 1,189 (594 + 595) | 1,339 (608 + 731) |

<sup>1</sup> Number of genes with at least one *cis*-eQTL (within 1kb of genes) and number of genes with only *trans*-eQTLS

We compared eQTLs mapped in females and males and asked whether the genetic control of gene expression by individual eQTLs is preserved in the two sexes. Consistent with the widespread prevalence of sexual dimorphism and sex by line interaction in gene expression, there are only 185 genes with at least one common eQTL in both sexes (Figure S14). The remaining genes contain either sex-specific eQTLs or do not vary genetically in the other sex (Figure S14).

To assess the fraction of total genetic variance explained by mapped eQTLs, we first identified eQTLs for each expression trait that are largely independent. To do this, we performed forward model selection to successively add eQTLs to an additive genetic model for each genetically variable gene expression trait, requiring that the conditional *P*-value of each added eQTL was smaller than 10^−5^. The number of eQTLs selected by the forward selection ranged from one to seven, with the majority of gene expression traits having one or two independent eQTLs (Figure 4A-B, Tables S10, S11). For most genes, the selected eQTLs explained a substantial fraction of genetic variance (Figure 4C-D).

**Figure 4:**
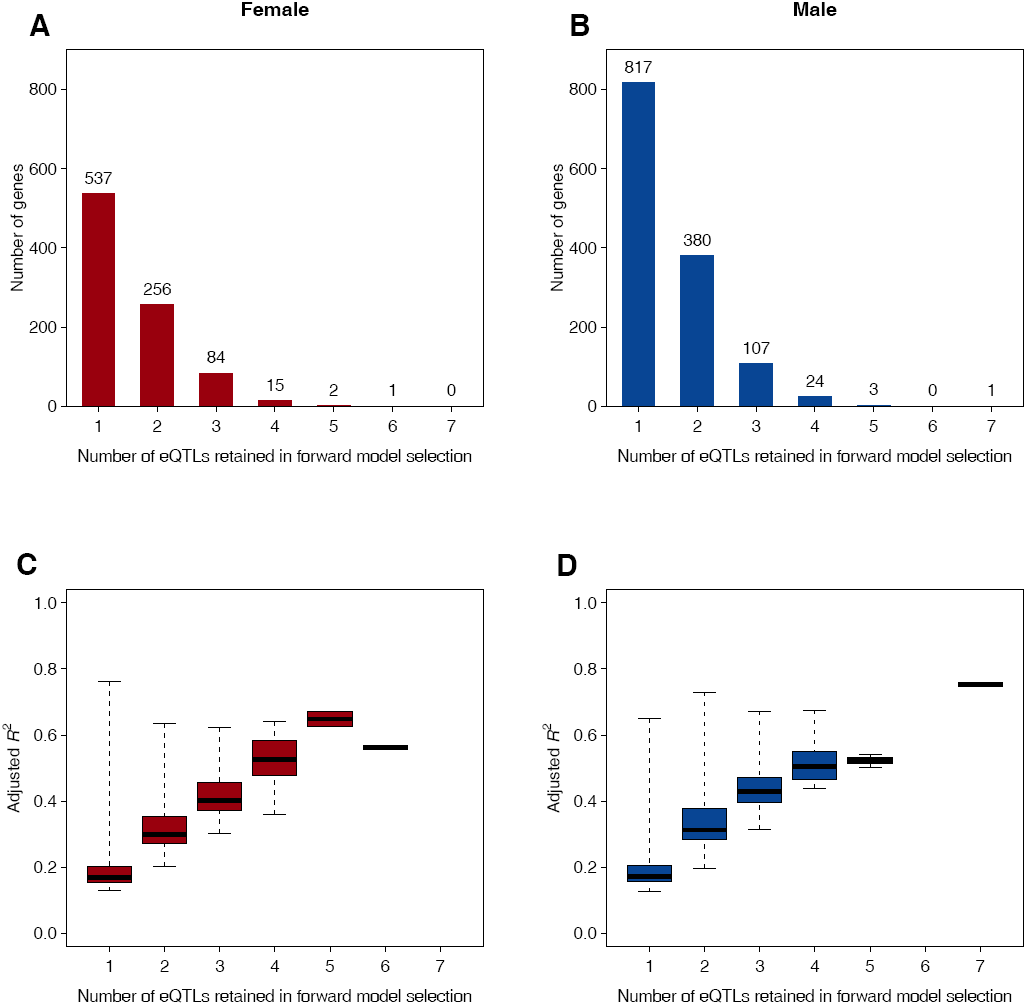
Variance in gene expression explained by independent eQTLs. (**A** and **B**) Distributions of the numbers of eQTLs retained in forward model selection. (A) Females.(B) Males. (**C** and **D**) Genetic variance explained by detected eQTLs (as measured by adjusted *R^2^)* versus the number of selected eQTLs. (C) Females. (D) Males.

Finally, we performed gene-based tests to search for groups of low frequency (MAF < 0.05) variants within 1kb of gene boundaries that collectively affect local gene expression. We used permutation to estimate the empirical FDR. At an FDR < 0.20, 626 (females) and 1,153 (males) genes are significantly associated with *cis*- low frequency variants (Tables S12, S13). Remarkably, 216 (females) and 408 (males) of these genes also contain common eQTLs in *cis*, accounting for more than 75% of all genes with a common *cis*-eQTL. This result suggests that mapping eQTLs with common frequencies also captures effects induced by rare variants collectively.

### QTLs associated with variance of expression

To search for variance eQTLs (veQTLs) for which lines carrying different alleles differ in their variance of expression within lines carrying the same allele, we performed a genome-wide scan for each gene expression trait using Levene’s test (Levene, 1960) for homogeneity of variance between two groups. At an FDR < 0.20, 925 and 412 genes in females and males contained at least one veQTL respectively (Table 2), among which 47 and 0 are NTRs (Tables S14, S15). The vast majority are *trans*-veQTLs (Table 2) and correspondingly, the strength of association between veQTLs and variance among lines within the same genotype class showed only weak concentration around TSS and TES (Figure S15).

**Table 2.** Number of genes with at least one significant veQTL at different FDR thresholds.

| Sex | FDR thresholds (*cis* + *trans*)<sup>1</sup> |  |  |  |
| --- | --- | --- | --- | --- |
|  | 0.05 | 0.10 | 0.15 | 0.20 |
| Female | 319 (6 + 313) | 544 (8 + 536) | 743 (9 + 734) | 925 (9 + 916) |
| Male | 162 (3 + 159) | 247 (6 + 241) | 353 (7 + 346) | 412 (7 + 405) |

<sup>1</sup> Number of genes with at least one *cis*-veQTL (within 1kb of genes) and number of genes with only *trans*-veQTLs

To obtain veQTLs that are independent from each other, we successively selected veQTLs from those that met the initial FDR thresholds. For each gene with more than one significant veQTL, we started with the most significant veQTL and scaled the variance of gene expression within the major and minor allele classes to unit variance while preserving their means. We then tested the next veQTL in the *P*-value ranked list of veQTLs using the scaled phenotype and continued this process until no veQTL could be added with a *P*-value smaller than 10^−5^. Similar to the mean eQTL analysis, this forward selection procedure also led to few veQTLs that independently controlled the variance of gene expression (Figure S15). Consistent with the observation that veQTLs were concentrated only weakly around genes (Figure S16), few genes with veQTLs contained *cis*-veQTLs (Table 2) after forward selection, a sharp contrast to eQTLs (Table 1).

Of the 941 genes in females and 1,339 genes in males whose expression was controlled by at least one eQTL, 248 and 107 respectively also had veQTLs. In total, 1,618 genes in females and 1,644 genes in males had at least one eQTL or veQTL, *i.e.* at least partially under the control of regulatory DNA variants. We could not assess whether genes with eQTLs are more likely to have veQTLs because the magnitude of variation between lines affects the power to detect both veQTLs and eQTLs. We further asked whether there were variants that control both the mean and variance in expression of the same genes. Among the 1,432 eQTL gene pairs in females and 2,029 in males retained in forward model selection, 16 and 6 were also significantly associated with the same genes as veQTLs respectively. Of these mean eQTLs that were also variance eQTLs, 1 and 0 were *cis* (< 1kb within genes) in females and males and the remaining were *trans.* On the other hand, among the 1,170 and 484 veQTL pairs in females and males, 24 and 15 were also significantly associated with the same genes as eQTLs, and 4 and 4 were *cis*, respectively in females and males. Moreover, only 37 of the 1,170 veQTLs in females and 28 of 484 in males showed significant association with the mean expression of any genes, suggesting that the variance-controlling effects of veQTLs were generally not due to their effects on changing the mean level of expression of other genes. Taken together, these results suggest that the genetic architectures for mean and variance of gene expression are largely independent.

### veQTLs are involved in epistatic interactions with *cis*-eQTLs

Because veQTLs can be emergent effects of underlying epistatic interactions for mean expression, we looked for variants that interact with veQTLs to epistatically affect gene expression. Because of the large number of possible epistatic pairs genome-wide, we limited the search to interactions between veQTLs and variants that are in *cis* (within 1kb) to the genes affected by the veQTLs. At an empirical FDR < 0.20, the vast majority (727 of 925 for females and 348 of 412 for males) of veQTLs for genes interacted with at least one *cis* variant (Figure S17). Moreover, among the 248 genes in females and 107 genes in males that had both eQTLs and veQTLs, 86 and 41 respectively had detectable interactions between the *cis* eQTLs and the veQTLs. For example, the expression of the Serine protease 12 *(Ser12)* gene in females was associated with a *cis* eQTL (Figure 5A) and a *trans* veQTL (Figure 5B), which interacted epistatically to change the mean of expression for individuals carrying the same allelic combinations (Figure 5C). The effect of the *cis* eQTL for *Ser12* therefore depended on the genotype of the *trans* veQTL (Figure 5D), which nevertheless was detected by ignoring the veQTL genotype in this specific case. However, many more *cis* variants have veQTL dependent effects that could not be detected by single marker regression (Figure 5E-H), highlighting the complexity and importance of context dependent effects in the genetic architecture of gene expression.

**Figure 5:**
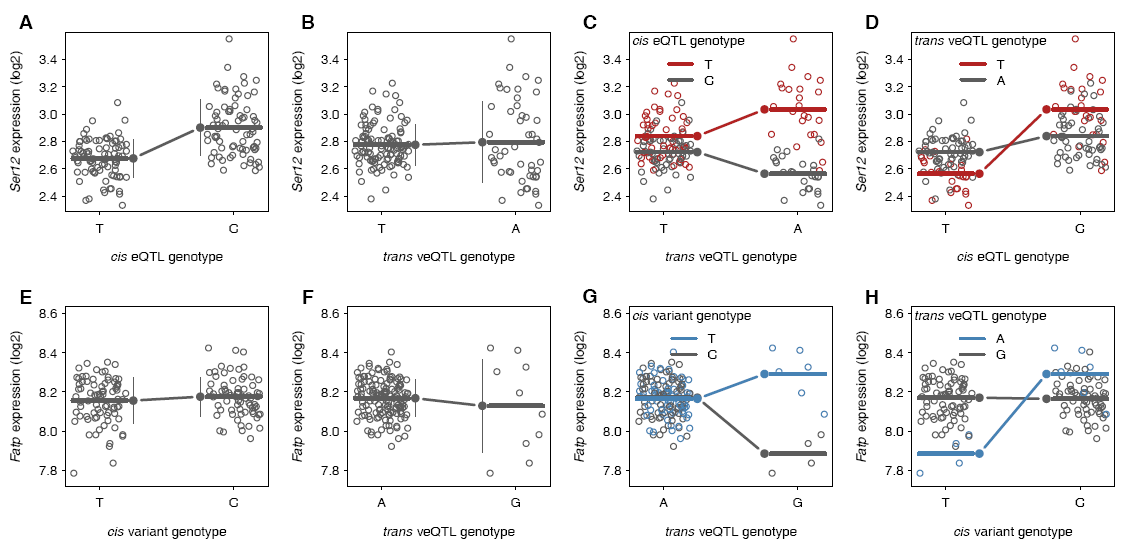
veQTLs are involved in epistatic interaction with *cis* variants. (**A**-**D**) Scatter plots of *Serl2* (*2L*:2250431..2251275) expression in females versus eQTL or veQTL genotypes. (A) The effect of a *cis* eQTL (*2L_2251218_SNP*) on the mean but not variance expression of individuals carrying the same genotypes. (B) The effect of a *trans* veQTL (*2L_11857529_SNP*) on the variance of but not the mean of expression of individuals carrying the same genotypes. (C) The effect of the *trans* veQTL on the mean expression is dependent on the *cis* eQTL genotype. (D) The effect of the *cis* eQTL on the mean expression is dependent on the *trans* veQTL genotype. (**E**-**F**) Scatter plots of *Fatp* (*2L*: 10510672..10517218) expression in males versus eQTL or veQTL genotypes. (E) No effect of a *cis* variant (*2L_10510716_SNP*) on the mean or variance of expression. (F) The effect of a *trans* veQTL (*3L_17881605_SNP*) on the variance of but not the mean of expression. (G) The effect of the *trans* veQTL on the mean expression is dependent on the *cis* variant genotype. (H) The effect of the *cis* variant on the mean expression is dependent on the *trans* veQTL genotype.

### Discussion

We have performed a comprehensive population-scale genetic characterization of the *D. melanogaster* transcriptome in a genetic reference population of sequenced, inbred, wild-derived lines. Similar to a previous study based on a subset of DGRP lines, we find that there is pervasive sexual dimorphism in mean gene expression and that a substantial fraction of the transcriptome is genetically variable (Ayroles et al., 2009; Massouras et al., 2012). In contrast to the previous studies, which utilized Affymetrix 3’ IVT microarrays, this analysis employed genome tiling microarrays. In this study we observed lower levels of genetic variance, higher within line variation, and correspondingly lower average heritabilities than observed previously. However, this decrease in precision was offset by our ability to assess the considerable contribution of NTRs to genetic variation in gene expression.

The abundances of genetically variable genes are not independent, but co-vary and form highly connected gene expression modules (Ayroles et al., 2009). These co-expression modules are not purely statistical constructs but are enriched for GO categories; and, reciprocally, genes in the same GO category tend to be genetically correlated. The highly genetically correlated transcriptome sets the stage for annotating genes for which there is no functional information using the ‘guilt by association’ principle, which is particularly useful for NTRs that have not been annotated previously. Several hundred of these NTRs were genetically variable and tend to correlate negatively with protein-coding genes. We functionally annotated many of the previously unknown NTRs based on their genetic correlations with gene expression of known genes. Despite their weak conservation and low expression levels, many NTRs may have biological functions based on their association with genes of known functions. Further characterization of these NTRs and their mechanism(s) of regulation of transcription is an exciting area for future investigation.

We performed GWA analyses to identify eQTLs for mean gene expression as well as for variance of expression in the DGRP. In both cases we used a stringent forward model selection procedure to avoid over-fitting QTLs. These analyses revealed that the genetic basis of transcriptional regulation is sex-specific, and largely independent for the mean and variance. Most transcripts had single eQTLs or veQTLs (a consequence of the model selection criteria), although 40% of mean expression traits had more than one eQTL and 15-23% of variance expression traits had more than one veQTL. Males had relatively more eQTLs and fewer veQTLs than females. At an FDR < 0.05, most eQTLs are in *cis*- to the gene whose expression they regulate, and typically map near transcription start and end sites, as has been shown previously in *D*. *melanogaster* and other species (Ronald et al. 2005; Stranger et al. 2007; Veyrieras et al. 2008; Massouras et al. 2012). The numbers of *trans*-eQTLs increase as the FDR threshold is lowered. In contrast, the majority of veQTLs are *trans*- to the gene for which they regulate variance in expression, and the fraction of *cis*-veQTLs remains low as the FDR threshold is lowered.

eQTLs in humans are enriched in *cis* regulatory elements such as DNase I hypersensitive sites, chromatin marks, and transcription factor binding sites (Brown et al. 2013). In contrast, little is known about the regulatory nature of veQTLs. It has been postulated that veQTLs could reflect underlying genetic (epistatic) or genotype by environment interactions (Brown et al. 2014). Here, we demonstrated that *trans*-veQTLs frequently interact epistatically with *cis*-variants to modulate gene expression levels (Figure 5, S17).

However, these interacting *cis*-variants are not the same as those affecting mean gene expression. The exact mechanisms are likely gene specific and remain to be studied.

The influences of sex and genetic interactions on gene expression fall into the broad framework of context-dependent effects, which provide the basis for dynamic gene expression programs during development and in response to different physical and social environments. Indeed, a substantial fraction of the *Drosophila* transcriptome is plastic and sensitive to changing environments (Zhou et al. 2012). However, the genetic basis of such plasticity is yet to be determined. The present study provides a baseline for further studies that investigate transcriptome diversity under various conditions.

In summary, the genetic architecture of *Drosophila* gene expression is complex and sex-specific, with pervasive genetic correlation between gene expression traits presumably caused in part by pleiotropy, and loci affecting both mean and variance in expression, the latter of which is frequently attributable to epistatic interactions (Figure 6). Epistatic interactions have also been implicated in the genetic architecture of complex traits (Huang et al. 2012; Mackay 2014). These complexities need to be incorporated into systems genetics models seeking to predict organismal level phenotypes for quantitative traits from gene expression data (Mackay et al. 2009). Further, our estimates of gene expression were from tiling arrays, which have a narrow dynamic range relative to digital gene expression estimates from RNA sequencing, and from entire flies at a single age and environmental condition. Further work is needed to assess to what extent these features of the genetic architecture of gene expression are robust or plastic in different tissues, developmental stages and social and physical environments.

**Figure 6:**
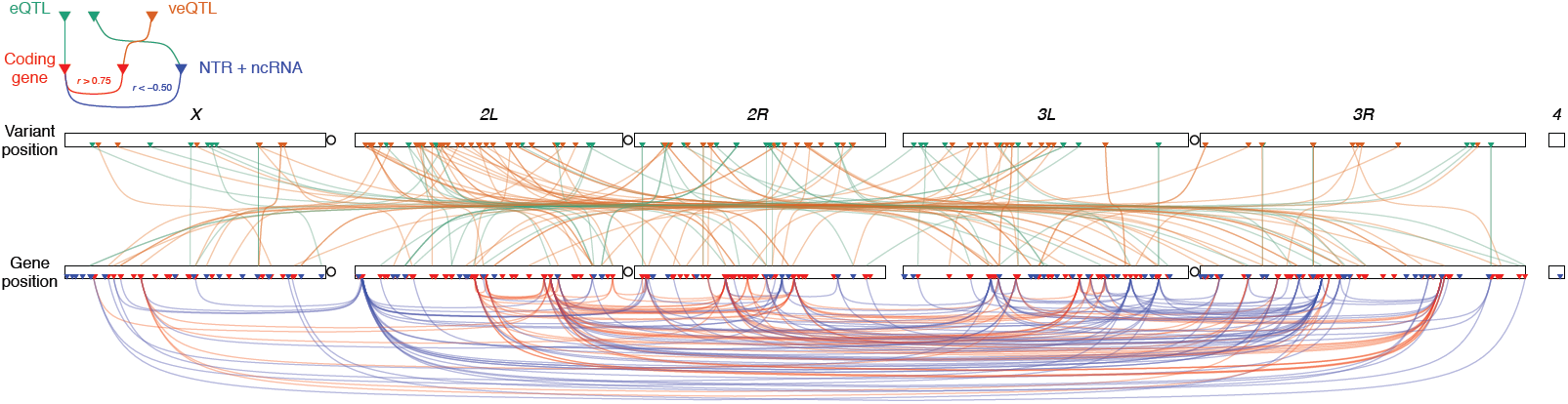
Architecture of genetic variation and genetic correlation in gene expression. The relationships between eQTL-gene, veQTL-gene, and gene-gene pairs are shown. Physical locations of DNA variants (chromosomes on top) and genes (chromosomes on bottom) are indicated by triangles, where red, brown, red, and blue triangles denote eQTL, veQTL, protein coding genes, and NTR or ncRNA, respectively. Green lines connect eQTLs and their associated genes, brown lines connect veQTLs and their associated genes, while red lines connect genes whose expression correlate at *r* > 0.75 and blue lines connect genes whose expression correlate at *r* < −0.5.

## Methods

### Drosophila lines

We used inbred lines of the *Drosophila melanogaster* Genetic Reference Panel (DGRP). These lines were established by 20 generations of full sib inbreeding from isofemale lines established from gravid females collected at the Raleigh, USA Farmer’s Market. Complete genome sequences of the DGRP lines have been obtained using the Illumina platform. SNPs, indels and other complex non-SNP variants have been genotyped using an integrated genotyping method (Huang et al. 2014).

### Fly husbandry and collection

All lines were reared under standard culture conditions (cornmeal-molasses-agar medium, 25°C, 60–75% relative humidity and a 12-hr light-dark cycle) at equal larval densities. For each line, we collected two replicates per sex for analysis of gene expression, consisting of 25 female flies or 40 male flies per replicate (∼ 25 mg each), for a total of 768 samples. Since it was not possible to collect all replicates from all lines simultaneously, we used a strict randomized experimental design for sample collection. We collected mated 3-5 day old flies between 1-3 pm. We transferred the flies into empty culture vials and froze them over ice supplemented with liquid nitrogen, and sexed the frozen flies. The samples were transferred to 2.0 ml nuclease-free microcentrifuge tubes (Ambion) and stored at −80°C until ready to process.

### RNA Extraction

The flies were homogenized with 1 ml of QIAzol lysis reagent (Qiagen) and two ¼ inch ceramic beads (MP Biomedical) using the TissueLyser (Qiagen) adjusted to a frequency of 15 Hz for 1 minute. Total RNA was extracted using the miRNeasy 96 kit (Qiagen) with on-column DNAse I digestion and following the spin technology protocol as outlined in the manufacturer’s manual. The RNA was eluted with 45 μl of RNAse-free water. The eluted samples contain total RNA including miRNAs and other small RNAs (≥18 nucleotides). Total RNA was quantified using a NanoDrop 8000 spectrophotometer (Thermo Scientific) and the concentrations of the RNA samples adjusted to 1 μg/μl for preparation of biotin-labeled double-stranded cDNA.

### RNA-Seq annotation of DGRP lines

We pooled 200 ng total RNA from each of 192 DGRP lines, separately for males and females. Poly(A)+ RNA-Seq libraries were prepared from each pool according to the Illumina TruSeq mRNA-Seq protocol, multiplexed, and sequenced by 100 bp paired-end in one lane of the HiSeq 2000 platform. Approximately 100 M fragments were sequenced for each of the male and female libraries. Sequence reads were mapped to the transcriptome (FlyBase annotation r5.49) and genome (FlyBase r5.49) using TopHat (version 2.0.8 with bowtie2-2.1.0, Trapnell et al. 2009), allowing a maximum edit distance of 6 bp. Gene models were assembled for male and female separately from the cDNA alignments using Cufflinks (version 2.0.2, Trapnell et al. 2010; Roberts et al. 2011) with the guide of the reference annotation. The transcript assemblies from males and females were merged and compared with the reference annotation to identify transcripts in previously unannotated intronic and intergenic **N**ovel **T**ranscribed **R**egions (NTRs).

### Preparation of whole transcript double-stranded cDNA

For each of the two replicates for each line and each sex, first strand cDNA was prepared from 7 μg of total RNA (1 μg/μl) with 1 μl of random primers (3 μg/μl) (Invitrogen) and incubating at 70°C (5 minutes) followed by 25°C (5 minutes) and 4°C (10 minutes). We added 5x first-strand buffer (4 μl; Invitrogen), 0.1 M dithiothreitol (2 μl; Invitrogen), 10 mM dNTP+dUTP (1 μl; Promega), RNase Inhibitor (1 μl; Invitrogen) and SuperScript II (4 μl; Invitrogen) and incubated the reactions in a thermal cycler (with heated lid) using the following program: 25°C / 10 minutes; 42°C / 90 minutes; 70°C / 10 minutes; 4°C / 10 minutes. Second-strand cDNA was synthesized by adding 17.5 mM MgCl_2_ (8 μl; Sigma), 10 mM dNTP+dUTP (1 μl; Promega), DNA Polymerase I (1.2 μl; Promega), RNase H (0.5 μl; Promega) and RNase-free water (9.3 μl; Ambion) to the first-strand cDNA reactions. The reactions were incubated in a thermal cycler at 16°C for 2 hours (without heated lid) followed by 75°C for 10 minutes (with heated lid) and 4°C for 10 minutes. Double-stranded cDNA was purified using the QIAquick 96 PCR kit (Qiagen) by following the manufacturer’s protocol except that buffer PN was used instead of buffer PM. The cDNA was eluted with 45 μl of RNAse-free water and quantified using a NanoDrop 8000 spectrophotometer (Thermo Scientific).

### Fragmentation and biotin-labeling of double-stranded cDNA

The double-stranded cDNA (7.5 μg) was fragmented with 4.8 μl 10 X fragmentation buffer (Affymetrix), 1.5 μl UDG (10 U/μl; Affymetrix), 2.25 μl APE 1 (100 U/μl; Affymetrix) and RNase-free water (up to 48 μl; Affymetrix) using a thermal cycler (with heated lid) and the following program: 37°C (1 hour), 93°C (2 minutes), 4°C (10 minutes). The fragmented double-stranded DNA (45 μl) was biotin-labeled by incubation with 12 μl of 5X TdT buffer (Affymetrix), 2 μl of 30 U/μl TdT (Affymetrix) and 1 μl of 5 mM DNA labeling reagent (Affymetrix) in a thermal cycler (with heated lid) using the following protocol: 37°C (1 hour), 70°C (10 minutes) and 4°C (10 minutes). Hybridization cocktail (164 μl) was added to 7 μg of fragmented and labeled double-stranded cDNA for hybridization to *Drosophila* 2.0R Tiling Arrays (Affymetrix). We randomized RNA extraction, labeling and hybridization across all samples.

### Quality control

We visualized the spatial distribution of probe intensities using the R package ‘Starr’ to identify technical artifacts on the arrays (e.g., salt rings from reagents). We also considered arrays to be outliers if the mean expression of probes on the array was ± two standard deviations of the sample mean from all arrays in the study; or if the variance of probe expression was ± two standard deviations from the sample mean variance of arrays in the study. We re-hybridized samples from all arrays with visible spatial artifacts and all outlier arrays to new arrays, using the same labeled samples used for the original arrays. Of the 192 lines that were initially hybridized to Affymetrix arrays, we retained 185 lines for analysis that have sequence data. Finally, within each sex, we removed replicates that contained excessive numbers of genes that were ± two standard deviations from the sample mean. A total of three replicate arrays (two female and one male replicate) were removed.

### Preprocessing of tiling array data

Raw intensities of tiling arrays were extracted from the .CEL files using the R package ‘AffyTiling’ and subjected to background correction on a per-array basis using functions modified from the ‘gcrma’ (version 2.30.0) package to work with tiling arrays. Briefly, non-specific binding affinities were calculated using 33,886 background probes on each array with varying degrees of GC content. The affinity information was then used to adjust for background hybridization for all *D. melanogaster* genomic probes on each array through a model-based approach (Wu et al. 2004). We mapped probes to the reference genome using BWA (version 0.6.2, Li and Durbin 2009) and removed probes that perfectly matched multiple genomic locations. Probes that fell entirely within non-overlapping constitutive exons as defined by the Flybase annotation (5.49) as well as NTRs discovered in the RNA-Seq annotation were retained. We further removed probes that overlapped with common (non-reference allele frequency > 0.05) variants in the DGRP Freeze 2.0 data (Huang et al. 2014). Background corrected intensities for the remaining 499,817 probes were quantile normalized (Bolstad et al. 2003) within each sex across arrays using the ‘limma’ (version 3.14.1) package. Expression for each gene was summarized using median polish.

### Quantitative genetics of gene expression

For each gene expression trait, we fitted a linear mixed model to partition variation in gene expression into the fixed effects of sex (S, sexual dimorphism in gene expression) and random effects of line (*L*, genetic variance) and the sex by line (*SL*, genetic variation in the magnitude of sex-dimorphism) interaction. The significance of sex effect was tested using a likelihood ratio test comparing the full model and a reduced model without the sex effect. The models were fitted using the ‘lme4’ package (version 0.999999-0) in R by maximum likelihood (ML). The significance of the sex by line variance was tested using an *F* test comparing the variance for the *SL* term and error variance. To estimate broad sense heritability (*H*^2^) for each gene expression trait in females and males separately, we fitted a linear mixed model with *L* as a random effect and estimated *H*^2^ as 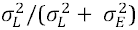 where 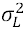 and 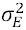 are, respectively, the between and within line variance components. The narrow sense heritability (*h*^2^) was estimated using a mixed linear model with *L* as a random effect and the covariance matrix determined by the genetic covariance among lines (Huang et al. 2014), using the ‘rrBLUP’ package (version 4.0) in R. The effect of *Wolbachia* and inversions were tested by a likelihood ratio test comparing the full model including *Wolbachia* infection status, inversion genotypes for *In(2L)t*, *In(2R)NS*, *In(3R)P*, *In(3R)K*, *In(3R)Mo*, and first ten principal components of the genotype matrix as fixed effects and *L* as a random effect, with a reduced nested model without the tested term. Principal components were obtained using the EIGENSTRAT software (Price et al. 2006) on LD pruned genotypes and excluding regions harboring the inversions.

### Gene set enrichment analysis

We performed gene set enrichment analysis (GSEA) on the list of genes ranked by their sex effect using a previously described procedure (Subramanian et al. 2005). We transformed *t* statistics to a signed correlation score 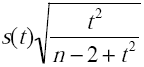, where *n* is the number of lines and *s*(*t*) indicates the sign of the t statistic. An empirical FDR was determined by permuting the sex label within each line 1,000 times and estimating the expected number of gene sets passing a certain threshold under the null hypothesis. Because the sex effect is large, unbalanced permutation can substantially bias the estimated sex effect. We removed one line from the data set to ensure that balanced permutation (the same number of females and males) can be properly performed. A similar GSEA was performed to annotate NTRs where the GSEA operated on the ranked list of annotated genes based on their correlation with the NTR.

### Mapping expression QTL (eQTL) for mean transcript abundance

eQTLs for mean gene expression were mapped using linear regression implemented in PLINK (Purcell et al. 2007), separately for males and females. The BLUP line means were first estimated using a mixed model adjusting for *Wolbachia*, inversions, and PCs, and then regressed on marker genotypes to obtain a *P*-value for each pair of markers and transcripts. To estimate the empirical FDR, we permuted line labels 100 times, retaining the correlation structure among the genes, and performed the same single marker regressions for the permuted phenotypes. The FDR was estimated by dividing the average number of significant markers meeting a certain threshold in the 100 permutations by the number of significant markers in the observed data set. To arrive at a model with independent associations, forward model selection was performed on significant markers. In each step, a marker with the smallest type III F test *P* value was added to the model until no marker could be added with a *P* < 10^−5^. Gene-based association tests were performed using the sequence kernel association test (SKAT; Wu et al. 2011) implemented in the ‘SKAT’ (version 0.95) package in R. The empirical FDR was determined using the same permuted data set and a similar procedure as described above for the marker-based tests.

### Mapping eQTL for variance of gene expression (veQTL) and epistasis

For each gene, veQTLs were mapped by testing for equal variance among the lines carrying the two alleles for each maker using Levene’s test. Empirical FDR was estimated by permutation as described above. To select for markers that independently control variance of gene expression, a forward selection procedure was performed on significant veQTLs. In each step, a maker with the smallest Levene’s test *P*-value was retained; after which the variance within each genotype class was scaled to unit variance while preserving the phenotypic mean. This process was repeated with the remaining markers until no marker could be added with a *P*-value smaller than 10^−5^. To identify *cis* variants that interact epistatically with veQTLs, the following model *y* = *μ* + *Mv* + *Mc* + *Mv:Mc* + *e* was fitted to each gene, where *y* is the adjusted gene expression, *μ* is an intercept, *Mv*, *Mc*, and *Mv:Mc* are the effects of the veQTL, *cis* variant, and their interaction respectively, and *e* is residual. This model was fitted for all pairs of veQTLs and all *cis* (within 1kb) variants of the gene. Significance of the interaction term was evaluated using an F test. Empirical FDR was calculated by permuting the gene expression and veQTL genotype together (thus a veQTL is still a veQTL after permutation) for 100 times and dividing the observed number of significant hits by the expected number of significant hits at variable thresholds.

## Data Access

The pooled RNA sequences from 192 DGRP lines have been deposited in Gene Expression Omnibus (GEO accession: GSE67505). All tiling array CEL files used in this study have been deposited at ArrayExpress (E-MTAB-3216).

## Acknowledgements

We thank Gunjan Arya, Julien Ayroles, Terry Campbell, Kultaran Chohan, Charlene Couch, Kyle Craver, Laura Duncan, Alden Hearn, George Khan, Faye Lawrence, Lenovia McCoy, Tatiana Morozova, Beth Ruedi, Yazmin Serrano-Negron, Shilpa Swarup, Crystal Tabor, Lavanya Turlapati, Allison Weber, Akihiko Yamamoto and Shanshan Zhou for technical assistance collecting samples for gene expression analysis. This work was supported by NIH grant R01 GM45146 to T.F.C.M., R.R.H.A. and E.A.S. and NIH grants R01 AA016560, R01 GM076083 and R01 GM59469 to T.F.C.M. and R.R.H.A. The authors declare they have no conflicts of interest.

## Supplementary Figure Legends

**Figure S1. Genomic characteristics of annotated genes and NTRs.** Violin plots showing (**A**) the distribution of transcript size, (**B**) GC content, (**C**) sequence conservation, and (**D**) variant density for annotated protein-coding genes, ncRNAs, and NTRs.

**Figure S2. Gene sets enriched for female- and male-biased transcripts.** Examples of significant gene sets enriched for (**A**) female-biased and (**B**) male-biased transcripts, from gene set enrichment analysis performed on genes ranked according to their sexual dimorphism in expression. The sexual dimorphism is measured by the correlation score, which is calculated as 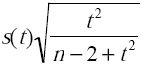, where *n* is the number of lines and *s*(*t*) indicates the sign of the *t* statistic for the difference in expression between males and females. Red and blue vertical bars indicate respectively the positions of female-biased and male-biased transcripts among the tested pathways on the ranked list. The purple line indicates the running enrichment score in the gene set enrichment analysis.

**Figure S3. Tissue-specific expression of sexually biased transcripts.** Tissue specificity of transcripts was calculated using the FlyAtlas gene expression profiles of 17 adult tissues as 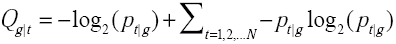, where 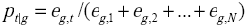 is the weighted expression of gene *g* in tissue *t*. *Q*_*g*|*t*_ is smaller when the expression of gene *g* is more specific in tissue *t*. In each tissue indicated, *Q*_*g*|*t*_ for all genes is plotted against the extent of sexual dimorphism as measured by the correlation score. The density of points on the plots is indicated by darkness of the color. The red line represents a smoothed curve computed by LOESS regression fit.

**Figure S4. Effects of *Wolbachia* infection on gene expression.** The effect of *Wolbachia* infection on gene expression was tested using a linear mixed model accounting for inversions and main PCs of the genotype matrix. The variance in gene expression explained by the presence or absence of *Wolbachia* is estimated as *p(1-p)w^2^*, where *p* is the proportion of lines infected by *Wolbachia* and *w* is the estimated mean change in gene expression upon infection. (**A** and **B**) Histograms of variance explained by *Wolbachia* infection. Colors indicate nominally significant transcripts. (A) Females. (B) Males. (**C** and **D**) Distributions of *P*-values for the *Wolbachia* effects, indicating a female-specific effect of *Wolbachia.* (C) Females. (D) Males. (**E** and **F**) Reproductive tissue specificity of transcripts with respect to their *Wolbachia* effect. (E) Specificity of ovary expression for genes expressed in females. (F) Specificity of testis expression for genes expressed in males. Genes that are down-regulated in *Wolbachia*-infected females show ovary-specific expression.

**Figure S5. Effects of inversions on gene expression.** The effects of major inversions on gene expression were tested using a linear mixed model accounting for inversions and main PCs of the genotype matrix. The variance explained by each inversion was estimated as 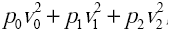 where *p*_*i*_ (*i* = 0,1,2) are the frequencies of lines that carry 0, 1, or 2 copies of the inverted karyotypes, and *v*_*i*_ (*i* = 0,1,2) are the estimated effects of each karyotype. This variance was divided by the total genetic variance to obtain the heritability explained by the inversions. (**A-J**) Plots depicting the heritability explained for transcripts, grouped according to their chromosomal regions for each of five major inversions. To test whether inverted regions are enriched for expression of genes affected by the inversion, the heritability of gene expression explained for genes in that region was compared with the remainder of the genome by a Wilcoxon test. The *P*-values from this test are indicated on each plot. (A) *In(2L)t*, females. (B) *In(2L)t*, males. (C) *In(2R)NS*, females. (D) *In(2R)NS*, males. (E) *In(3R)P*, females. (F) *In(3R)P*, males. (G) *In(3R)K*, females. (H) *In(3R)K*, males. (I) *In(3R)Mo*, females. (J) *In(3R)Mo*, males.

**Figure S6. Examples of enrichment of biological pathways in MMC modules.** The top three panels (red font) are for genetically correlated transcriptional modules in females, and the bottom four panels (blue font) are for male transcriptional modules. In each panel, the color intensity of the text (GO name) represents different levels of FDR according to the scale bar on the right. The size of the text represents the factor of enrichment with the scale for no enrichment (factor =1) indicated in the top left corner of each panel.

**Figure S7. Genes within the same biological pathways have highly correlated expression in females.** The box plots show the average connectivity (absolute correlation coefficient) within each GO pathway. The blue box depicts the connectivity among genes that have no known GO association.

**Figure S8. Genes within the same biological pathways have highly correlated expression in males.** The box plots show the average connectivity (absolute correlation coefficient) within each GO pathway. The blue box depicts the connectivity among genes that have no known GO association.

**Figure S9. *Wolbachia* infection has a minimal effect on genetic correlation of gene expression.** (**A** and **B**) Distributions of the difference between correlation coefficients between all pairs of genetically variable transcripts calculated with or without adjusting for the effect of *Wolbachia* infection on gene expression. (A) Females. (B) Males.

**Figure S10. NTRs are negative regulators.** (**A** and **B**) Distributions of correlation coefficients between protein-coding genes, between NTRs, and between NTRs and protein-coding genes, all within the same modules. (A) Females. (B) Males.

**Figure S11. Distance between NTRs and protein coding genes.** Distance between NTRs and protein-coding genes whose genetic correlation is greater than 0.25 within the same module or less than 0.25 in females (A) and males (B).

**Figure S12. Functional annotation of NTRs.** Annotation by association of NTRs with GO/KEGG pathways. The numbers of NTRs associated with GO/KEGG pathways are plotted for females (top panel) and males (bottom panel).

**Figure S13. Strength of association between local variants and gene expression.** (**A** and **B**) The plots depict the strength of association (−log_10_*P*) between DNA variants and gene expression (*y*-axis) against the distance relative to transcription start sites (TSS) or transcription end sites (TES). The figures show both the 95% quantile and median of – log_10_*P* for variants within each non-overlapping 200bp window, plotted against the midpoint of the window. Negative and positive distances indicate, respectively, positions upstream and downstream of the direction of transcription. The analysis is shown for gene expression traits with (FDR < 0.20) and without (FDR ≥ 0.2) eQTL. (A) Females. (B) Males.

**Figure S14. Sex-specific eQTLs.** Boxes on the left and right indicate the numbers of eQTL-gene pairs in females and males respectively. Ribbons (with the corresponding numbers of eQTL-gene pairs or genes) connect portions of the boxes that share the same eQTL-gene pairs or genes. The purple ribbon indicates shared eQTL-gene pairs in both sexes. The light red ribbon indicates eQTL-gene pairs that are significant only in females but not in males where the genes are also genetically variable. The light blue ribbon indicates eQTL-gene pairs that are significant only in males but not in females where the genes are also genetically variable. Grey ribbons indicate eQTL-gene pairs significant in one sex but not the other where the genes are not genetically variable.

**Figure S15. Strength of association between local variants and variance of gene expression.** (**A** and **B**) The plots depict the strength of association (−log_10_*P*) between DNA variants and variance of gene expression (*y*-axis) against the distance relative to transcription start sites (TSS) or transcription end sites (TES). The figures show both the 95% quantile and median of −log_10_*P* for variants within each non-overlapping 200bp window, plotted against the midpoint of the window. Negative and positive distances indicate, respectively, positions upstream and downstream of the direction of transcription. The analysis is shown for gene expression traits with (FDR < 0.20) and without (FDR ≥ 0.2) veQTL. (A) Females. (B) Males.

**Figure S16. veQTLs retained in model selection.** (**A** and **B**) Distributions of the numbers of veQTLs retained in forward model selection. (A) Females. (B) Males.

**Figure S17. veQTLs are involved in epistatic interactions with *cis* variants.** (**A** and **B**) Epistatic interactions between veQTLs and *cis* variants. (A) Females. (B) Males. Each veQTL and *cis* variant pair is connected by a light line. Interactions between *cis* veQTL and *cis* variants are highlighted by dark lines.

## References

Ayroles JF, Carbone MA, Stone EA, Jordan KW, Lyman RF, Magwire MM, Rollmann SM, Duncan LH, Lawrence F, Anholt RRH, et al. 2009. Systems genetics of complex traits in *Drosophila melanogaster*. Nat Genet 41: 299–307.

Ayroles JF, Laflamme BA, Stone EA, Wolfner MF, Mackay TF. 2011. Functional genome annotation of Drosophila seminal fluid proteins using transcriptional genetic networks. Genet Res 93: 387–395.

Brem RB, Yvert G, Clinton R, Kruglyak L. 2002. Genetic dissection of transcriptional regulation in budding yeast. Science 296: 752–755.

Brown AA, Buil A, Viñuela A, Lappalainen T, Zheng HF, Richards JB, Small KS, Spector TD, Dermitzakis ET, Durbin R. 2014. Genetic interactions affecting human gene expression identified by variance association mapping. eLife 3: e01381.

Brown CD, Mangravite LM, Engelhardt BE. 2013. Integrative modeling of eQTLs and cis-regulatory elements suggests mechanisms underlying cell type specificity of eQTLs. PLoS Genet, 9: e1003649.

Bolstad BM, Irizarry RA, Astrand M, Speed TP. 2003. A comparison of normalization methods for high density oligonucleotide array data based on variance and bias. Bioinformatics 19: 185–193.

Cheung VG, Conlin LK, Weber TM, Arcaro M, Jen KY, Morley M, Spielman RS. 2003. Natural variation in human gene expression assessed in lymphoblastoid cells. Nat Genet 33: 422–425.

Dinger ME, Amaral PP, Mercer TR, Mattick JS. 2009. Pervasive transcription of the eukaryotic genome: functional indices and conceptual implications. Brief Funct Genomic Proteomic 8: 407–423.

Djebali S, Davis CA, Merkel A, Dobin A, Lassmann T, Mortazavi A, Tanzer A, Lagarde J, Lin W, Schlesinger F. et al. 2012. Landscape of transcription in human cells. Nature 489: 101–108.

Flint J, Mackay TFC. 2009. Genetic architecture of quantitative traits in mice, flies and humans. Genome Res 19: 723–733.

Graveley BR, Brooks AN, Carlson JW, Duff MO, Landolin JM, Yang L, Artieri CG, van Baren MJ, Boley N, Booth BW, et al. 2011. The developmental transcriptome of *Drosophila melanogaster*. Nature 471: 473–479.

Hill WG, Goddard ME, Visscher PM. 2008. Data and theory point to mainly additive genetic variance for complex traits. PLoS Genet. 4: e1000008.

Huang W, Massouras A, Inoue Y, Peiffer J, Rámia M, Tarone A, Turlapati L, Zichner T, Zhu D, Lyman R, et al. 2014. Natural variation in genome architecture among 205 *Drosophila melanogaster* Genetic Reference Panel lines. Genome Res 24: 1193–1208.

Huang W, Richards S, Carbone MA, Zhu D, Anholt RRH, Ayroles JF, Duncan L, Jordan KW, Lawrence F, Magwire MM, et al. 2012. Epistasis dominates the genetic architecture of *Drosophila* quantitative traits. Proc Natl Acad Sci 109: 15553–15559.

Hulse AM, Cai JJ. 2013. Genetic variants contribute to gene expression variability in humans. Genetics 193: 95–108.

Lee JT. 2012. Epigenetic regulation by long noncoding RNAs. Science 338: 1435–1439.

Li H, Durbin R. 2009. Fast and accurate short read alignment with Burrows-Wheeler Transform. Bioinformatics 25: 1754–1760.

Levene H. 1960. Robust testes for equality of variances. In Contributions to Probability and Statistics: Essays in Honor of Harold Hotelling, (ed. Olkin I, et al.), pp. 278–292. Stanford University Press, Palo Alto, CA

Mackay TFC. 2014. Epistasis and quantitative traits: using model organisms to study gene-gene interactions. Nat Rev Genet 15: 22–33.

Mackay TFC, Richards S, Stone EA, Barbadilla A, Ayroles JF, Zhu D, Casillas S, Han Y, Magwire MM, Cridland JM, et al. 2012. The *Drosophila melanogaster* Genetic Reference Panel. Nature 482: 173–178.

Mackay TFC, Stone EA, Ayroles JF. 2009. The genetics of quantitative traits: challenges and prospects. Nat Rev Genet 10, 565–577.

Manolio TA, Collions FS, Cox NJ, Goldstein DB, Hindorff LA, Hunter DJ, McCarthy MI, Ramos EM, Cardon LR, Chakravarti A, et al. 2009. Finding the missing heritability of complex diseases. Nature 461: 747–753.

Massouras A, Waszak SM, Albarca-Aguilera M, Hens K, Holcombe W, Ayroles JF, Dermitzakis ET, Stone EA, Jensen JD, Mackay TFC, et al. 2012. Genomic variation and its impact on gene expression in *Drosophila melanogaster*. PLoS Genet 8: e1003055.

Nicolae DL, Gamazon E, Zhang W, Duan S, Dolan ME, Cox NJ. 2010. Trait-associated SNPs are more likely to be eQTLs: Annotation to enhance discovery from GWAS. PLoS Genet 6: e1000888.

Ober U, Ayroles JF, Stone EA, Richards S, Zhu D, Stricker C, Gianola D, Schlather M, Mackay TFC, Simianer H. 2012. Using whole-genome sequence data to predict quantitative trait phenotypes in *Drosophila melanogaster*. PLoS Genet 8: e1002685.

Price AL, Patterson NJ, Plenge RM, Weinblatt ME, Shadick NA, Reich D. 2006. Principal components analysis corrects for stratification in genome-wide association studies. Nature Genet 38: 904–909.

Parisi M, Nuttall R, Edwards P, Minor J, Naiman D, Lu J, Doctolero M, Vainer M, Chan C, Malley J. 2004. A survey of ovary-, testis-, and soma-biased gene expression in *Drosophila melanogaster* adults. Genome Biol 5: R40.

Purcell S, Neale B, Todd-Brown K, Thomas L, Ferreira MA, Bender D, Maller J, Sklar P, de Bakker PIW, Daly MJ et al. 2007. PLINK: a tool set for whole-genome association and population-based linkage analyses. AmJ Hum Genet 81: 559–575.

Ranz JM, Castillo-Davis CI, Meiklejohn CD, Hartl DL. 2003. Sex-dependent gene expression and evolution of the *Drosophila* transcriptome. Science 300: 1742–1745.

Roberts A, Pimentel H, Trapnell C, Pachter L. 2011 Identification of novel transcripts in annotated genomes using RNA-Seq. Bioinformatics 27: 2325–2329.

Rönnegård L, Valdar W. 2011. Detecting major genetic loci controlling phenotypic variability in experimental crosses. Genetics 188: 435–447.

Schadt, EE, Monks, SA, Drake, TA, Lusis, AJ, Che, N, Colinayo, V, Ruff TG, Milligan SB, Lamb JR, Cavet G, et al. 2003. Genetics of gene expression surveyed in maize, mouse and man. Nature, 422, 297–302.

Shen X., Pettersson M., Ronnegard L, & Carlborg Ö. 2012. Inheritance beyond plain heritability: variance-controlling genes in *Arabidopsis thaliana*. PLoS Genet, 8: e1002839.

Stone EA, Ayroles JF. 2009. Modulated modularity clustering as an exploratory tool for functional genomic inference. PLoS Genet 5: e1000479.

Subramanian A, Tamayo P, Mootha VK, Mukherjee S, Ebert BL, Gillette MA, Paulovich A, Pomeroy SL, Golub TR, Lander ES et al. 2005. Gene set enrichment analysis: a knowledge-based approach for interpreting genome-wide expression profiles. Proc Natl Acad Sci 102: 15545–15550.

Swarup S, Huang W, Mackay TFC, Anholt RRH. 2013. Analysis of natural variation reveals neurogenetic networks for *Drosophila* olfactory behavior. Proc Natl Acad Sci 110: 1017–1022.

Trapnell C, Pachter L, Salzberg SL. 2009. TopHat: discovering splice junctions with RNA-Seq. Bioinformatics. 25: 1105–1111.

Trapnell C, Williams BA, Pertea G, Mortazavi A, Kwan G, van Baren MJ, Salzberg SL, Wold BJ, Pachter L. 2010. Transcript assembly and quantification by RNA-Seq reveals unannotated transcripts and isoform switching during cell differentiation. Nature Biotech 28: 511–515.

Wu M, Lee S, Cai T, Li Y, Boehnke M, Lin X. 2011. Rare-variant association testing for sequencing data with the sequence kernel association test. Am J Hum Genet. 89: 82–93.

Wu Z, R.A. Irizarry, Gentleman R, Martinez-Murillo F, Spencer F. A model-based background adjustment for oligonucleotide expression arrays. J Amer Stat Assoc. 99, 909–917 (2004).

Yang J, Benyamin B, McEvoy BP, Gordon S, Henders AK, Nyholt DR, Madden PA, Heath AC, Martin NG, Montgomery GW, et al. 2010. Common SNPs explain a large proportion of the heritability for human height. Nat Genet 42: 565–569.

Yang J, Loos RJ, Powell JE, Medland SE, Speliotes EK, Chasman DI, Rose LM, Thorleifsson G, Steinthorsdottir V, Mägi R, Waite L, et al. 2012. FTO genotype is associated with phenotypic variability of body mass index. Nature 490: 267–272.

Zhou S, Campbell TG, Stone EA, Mackay TFC, Anholt RRH. 2012. Phenotypic plasticity of the *Drosophila* Transcripto me. PLoS Genet 8: e1002593.

